# Skeletal muscle mitochondrial respiration in a model of age-related osteoarthritis is impaired after dietary rapamycin

**DOI:** 10.1101/2021.06.11.448123

**Authors:** Christian J. Elliehausen, Dennis M. Minton, Alexander D. Nichol, Adam R. Konopka

## Abstract

A decline in skeletal muscle mitochondrial function is associated with the loss of skeletal muscle size and function during knee osteoarthritis (OA). We have recently reported that the 12-weeks of dietary rapamycin (Rap, 14ppm), with or without metformin (Met, 1000ppm), increased plasma glucose and OA severity in male Dunkin Hartley (DH) guinea pigs, a model of naturally occurring, age-related OA. The purpose of the current study was to determine if increased OA severity after dietary Rap and Rap+Met was accompanied by impaired skeletal muscle mitochondrial function. Mitochondrial respiration and hydrogen peroxide (H_2_O_2_) emissions were evaluated in permeabilized muscle fibers via high-resolution respirometry and fluorometry using either a saturating bolus or titration of ADP. Rap and Rap+Met decreased complex I (CI)-linked respiration and increased ADP sensitivity, consistent with previous findings in patients with end-stage OA. Rap also tended to decrease mitochondrial H_2_O_2_ emissions, however, this was no longer apparent after normalizing to respiration. The decrease in CI-linked respiration was accompanied with lower CI protein abundance. This is the first inquiry into how lifespan extending treatments Rap and Rap+Met can influence skeletal muscle mitochondria in a model of age-related OA. Collectively, our data suggest that Rap with or without Met inhibits CI-linked capacity and increases ADP sensitivity in DH guinea pigs that have greater OA severity.

## INTRODUCTION

Osteoarthritis (OA) is amongst the top 10 diseases that limit human healthspan and lifespan (Murray et al., 2013). However, there are currently no disease modifying therapies for OA. Age is the greatest risk factor for naturally occurring, primary OA. OA is the leading cause of disability in older adults affecting 250 million people worldwide and creating a significant individual and socioeconomic burden (Hunter et al., 2014). OA is characterized as a disease of the whole-joint, including articular cartilage, subchondral bone and periarticular skeletal muscle (Loeser et al., 2012). The loss of skeletal muscle size (Fink et al., 2007; Serrão et al., 2015), function (Øiestad et al., 2015; Thorstensson et al., 2004), and quality (Kumar et al., 2014; Noehren et al., 2018) are features of OA that contributes to joint pain, physical inactivity, and comorbidities. Although it is difficult to dissociate the loss of muscle size and function with OA as a result of muscle disuse, there is evidence to suggest that loss of quadriceps muscle strength may occur prior to and be predictive of incident radiographic and symptomatic knee OA (Segal & Glass, 2011; Slemenda et al., 1997).

The age-related decline of skeletal muscle mitochondrial function is associated with impaired skeletal muscle contractile function (Gonzalez-Freire et al., 2018), walking speed (Coen et al., 2013), and fatigability (Santanasto et al., 2015). Older adults with high fatigability also had lower mitochondrial respiration compared to older adults with low fatigability. Nearly 50% of older individuals with high fatigability were diagnosed with OA (Santanasto et al., 2015). Consistent with these findings, patients with end-stage OA about to receive knee arthroplasty had lower skeletal muscle mitochondrial content and maximal complex IV activity compared to young and age-matched non-OA controls (Safdar et al., 2010). Additionally, OA patients have impaired redox homeostasis as evident by decreased activity of the antioxidant MnSOD and increased oxidative damage (Safdar et al., 2010).

The onset of naturally occurring (primary) OA in humans is insidious and unpredictable which limits the study of OA to end-stage disease and makes it difficult to study the cellular processes that are involved in the onset, early progression, and treatment of OA. To circumvent this limitation, we use the Dunkin Hartley (DH) guinea pig model of primary OA. The DH guinea pig predictably develops naturally occurring OA by 5 months of age and demonstrates an age-associated progression to moderate OA by 8-9 months of age (Bendele & Hulman, 1988; Radakovich et al., 2018). The DH guinea pig highly resembles the OA pathology observed in humans as evident by cartilage and bone lesions that typically originate in the medial tibia and spread laterally with age (Huebner & Kraus, 2006; Kapadia et al., 2000). In addition to OA, the DH guinea pig displays a number of recognized age-related changes in human skeletal muscle including a decline in skeletal muscle density, muscle fiber size and mitochondrial protein synthesis rates (Musci et al., 2020). These characteristics make the DH guinea pig a desirable model to study the onset, progression and treatment of primary, age-related OA.

The FDA approved drug rapamycin (Rap) can extend lifespan in mice (Harrison et al., 2009) and when metformin is added to Rap (Rap+Met) there may be an even greater effect on lifespan extension (Strong et al., 2016). Treatments that extend lifespan are also intended to extend healthspan, the time spent without chronic age-related disease and disability. However, the therapeutic potential of Rap and Met for specific age-related diseases is incompletely understood. Despite previous work showing protective effects of Rap or Met on injury induced OA in young mice, we have recently shown that dietary Rap and Rap+Met treatment increased plasma glucose and worsened age-related OA severity in DH guinea pigs (Minton et al., 2021). Based on these findings, we sought to determine if the detrimental effects of Rap and Rap+Met on OA pathology and glucose homeostasis would be accompanied by impaired skeletal muscle mitochondrial bioenergetics.

## METHODS

### Animal Use

All animal procedures were approved by the Institutional Animal Care and Use Committee at the University of Illinois Urbana-Champaign. Data collection and analysis were completed at University of Wisconsin-Madison and William S. Middleton Memorial Veterans Hospital. The details of the experimental design and animal husbandry have been previously described in (Minton et al., 2021). In brief, Dunkin Hartley Guinea Pigs were obtained from Charles River and during acclimation provided a standard chow diet (Envigo 2040) fortified with vitamin C (1050 ppm) and Vitamin D (1.5 IU/kg) *ad libitum*. At 5 months of age, DH guinea pigs were then randomized to continue the control diet (n=8), or standard chow enriched with encapsulated Rapamycin (14 ppm, n=8) or the combination of encapsulated Rapamycin+Metformin (14 ppm, 1000 ppm, n=7) for 12-weeks. Microencapsulated Rapamycin (Rapamycin holdings) and Metformin HCl (AK Scientific I506) were compounded into the diet by Envigo. Guinea pigs provided experimental diets enriched with Rap or the combination of Rap+Met were allowed continued *ad libitum* access to food and water. Food provided to the control group was calculated to match the food consumption of the experimental diet to minimize the influence of food intake on dependent variables.

### Tissue Collection

Following 3 months of treatment, two animals were sacrificed daily. Food was removed from cages 2-3 hours prior to euthanasia. Animals were anesthetized in a chamber containing 5% isoflurane and maintained using a mask with 1.5-3% isoflurane. Once reflexively unresponsive, the thoracic cavity was opened and whole blood was collected via cardiac venipuncture in K2EDTA coated tubes. The heart was then excised. The right soleus was removed from the hind limb and immediately placed in ice-cold BIOPS solution (2.77 mM CaK_2_-EGTA, 7.23 mM K_2_-EGTA, 20 mM imidazole, 20 mM taurine, 50 mM K-MES, 0.5 mM dithiothreitol, 6.56 mM MgCl_2_, 5.77 mM ATP, and 15 mM phosphocreatine, pH 7.1). The left soleus was frozen in liquid nitrogen and stored at −80C until processing.

### High-resolution respirometry and fluorometry

High-resolution respirometry and fluorometry measurements were performed in duplicate using two chambers of the Oxygraph-2k (O2K; OROBOROS INSTRUMENTS, Innsbruck, Austria). Air calibration and instrument background oxygen flux were performed before each experiment. After the addition of the permeabilized fibers, the O2K was calibrated to measure to measure H_2_O_2_ emissions with three injections of H_2_O_2_ (40μM). Temperature within the chambers was maintained at 37° C and oxygen concentrations were kept within the range of 350 - 250 μmol/mL. Reoxygenation was performed when oxygen concentrations approached 250 μmol/mL to ensure oxygen availability was not a limiting factor. Data are presented as the average between duplicates and respiration rates (O_2_ flux) and H_2_O_2_ emissions were expressed relative to tissue wet weight (pmol/s/mg).

### Tissue Preparation

Consistent with muscle from patients with knee OA, muscle from the Hartley guinea pig contained a visible prevalence of non-contractile tissue (Noehren et al., 2018). During mechanical permeabilization, connective and adipose tissue were removed. Fibers were then chemically permeabilized in BIOPS with saponin (50μg/mL) for 30 minutes. Permeabilized muscle fibers were rinsed with MiR05 (0.5 mM EGTA, 3 mM MgCl_2_, 60 mM K-lactobionate, 20 mM taurine, 10 mM KH_2_PO_4_, 20 mM HEPES, 110 mM sucrose, 1 g/l BSA essentially fatty acid free, pH 7.1) plus 25μM blebbistatin, a myosin II-specific inhibitor to prevent muscle fiber contraction (Perry et al., 2011). Fibers were then blotted on filter paper to remove excess buffer and weighed on a microscale (Mettler Toledo X105). Fibers bundles (1.5-2.5 mg) were then placed into the chambers containing MiR05 plus 12.5μM blebbistatin.

### Mitochondrial Bioenergetics

Two different protocols were used to assess mitochondrial function. The first protocol was supported by complex-I linked substrates, pyruvate (5mM), glutamate (10mM) and malate (0.5mM) to evaluate complex I (CI)-linked LEAK (CI_L_) respiration followed by a bolus of ADP (5mM) to stimulate maximal complex I-linked OXPHOS (CI_P_). Subsequent additions included cytochrome c (10mM) to test mitochondrial membrane integrity, succinate (10mM) for complex I plus II supported OXPHOS (CI+II_P_), and Carbonyl cyanide m-chlorophenyl hydrazone (CCCP) (0.25-0.75 μM) to stimulate maximal uncoupled electron transport system (ETS; CI+II_E_) capacity. Next, the complex-I inhibitor rotenone (0.5μM) was added so that the remaining respiration was reflective of CII_E_. Mitochondrial respiration was stopped by the complex III inhibitor antimycin A (2.5μM) to measure residual oxygen consumption (ROX). SUIT 1 was used to measure respiration in 4 animals per group.

We next utilized an ADP titration protocol to evaluate mitochondrial respiration and H_2_O_2_ emissions in a different set of 4 animals per group. This protocol was also supported by pyruvate (5mM), glutamate (10mM) and malate (0.5mM) to stimulate LEAK respiration and evaluate CI-linked H_2_O_2_ emissions. Subsequently, ADP was serially injected to reach concentrations of 0.125, 0.25, 0.56, 1.2, 2.4, 5.6, and 11.8 mM followed by sequential addition of cytochrome c, succinate, and CCCP to determine maximal CI+II_P_ and CI+II_E_ as previously performed (Konopka et al., 2017, 2019). Using Michaelis-Menten kinetics, V_max_ and the apparent Km for ADP were calculated. We also determined the area under the curve (AUC) for respiration and H_2_O_2_ emissions during titration of ADP. For both protocols, the ratio of maximal OXPHOS to maximal uncoupled ETS capacity (P/E) was used to gain insight into intrinsic mitochondrial function independent of changes in mitochondrial abundance. We also expressed H_2_O_2_ emissions relative to respiration as an indicator of electron leak.

### Western Blotting

Soleus and liver tissue were homogenized in liquid nitrogen cooled mortar and pestle followed by bead homogenization in RIPA lysis buffer (150mM NaCl, 0.1mM EDTA, 50mM Tris, 0.1% wt/vol deoxycholate, 0.1% wt/vol SDS, 1% vol/vol Triton X-100) with 1X Halt Protease and Phosphatase Inhibitor Cocktail (Thermofisher Scientific: 78442) as previously described (Minton et al., 2021) 10 μg of protein was loaded for OXPHOS proteins using a cocktail of subunits from the 5 electron transport system complexes, CI-NDUFB8, CII-SDH8, CIII-UQCRC2, CIV-MTCO1, CV-ATP5A (Abcam, MS604). 20 μg of soleus and liver protein loaded for all other proteins. Samples prepared in reducing conditions with β-mercaptoethanol in 2x Laemmli Sample Buffer (BioRad 1610737) and heated at 95C. The samples for OXPHOS blots were heated at 50C to preserve Complex I, II, IV proteins which are sensitive to high heat. Proteins were separated on 4-15% TGX Precast Gels (BioRad, 4561083) and transferred to PVDF membranes (Invitrogen). Membranes were blocked in TBS-Tween 20 (0.1%) (TBST) with 5% BSA and then incubated overnight with primary antibodies for OXPHOS (Abcam MS604), P-RPS6 Ser235/236 (Cell Signaling, 4858), RPS6 (Cell Signaling, 2217), P-AKT S473 (Cell Signaling, 4060), AKT (Cell Signaling, 4685) and B-Actin (Abcam Ab6276) at 1:500, 1:2000, 1:1000, and 1:5000 respectively. Despite reactivity with other guinea pig tissues, we were not able to detect AMPK (Cell Signaling, 2532) and p-AMPK (Cell Signaling, 50081) in DH guinea pig skeletal muscle therefore we probed liver tissue. OXPHOS was diluted in 1% Nonfat milk/PBS following manufacturer recommendation and all others in 5% BSA/TBST. Membranes were washed with TBST between incubations. Membranes were then incubated with IgG HRP conjugated Anti-Mouse (Abcam, 6728) and Anti-Rabbit (Cell Signaling, 7074) secondary antibodies at 1:10,000. Membranes exposed to SuperSignal Pico Substrate (Fisher, PI34095) and imaged with BioRad ChemiDox XRS+ or UVP Biospectrum 500. Membranes were stripped with Restore Stripping Buffer following manufacturer recommendations (ThermoFisher Scientific 21059). Densitometric calculations determined using Image Lab (BioRad) or VisionWorks (Analytikjena).

### Statistics

Data are presented as mean ± standard error of the mean (SEM). Significance was set at P < 0.05. Michaelis-Menten kinetics were used to determine the estimated apparent K_m_ and maximal respiration (V_max_) in Prism 9.0 (GraphPad Software, Inc., La Jolla, CA, USA). One-way ANOVA was used. When significant main effects were present a two-stage linear step-up procedure of Benjamini, Krieger and Yekutieli post-hoc test was chosen to account for multiple comparisons. A priori, our primary aim was to test the effects of each treatment compared to control and not to determine differences between treatments.

## RESULTS

### DH Guinea Pig Characteristics

We have previously published the characteristics and OA score from the same cohort of DH guinea pigs (Minton et al., 2021). We summarize those data in Table 1 for ease of readership. Briefly, Rap and Rap+Met animals weighed 15 and 22% (P<0.05) less than control animals. Rap and Rap+Met treated animals had increased fasting plasma glucose by 64 and 39% (P<0.05). OA severity measured by medial tibia OARSI score was nearly 2-fold greater in Rap and Rap+Met compared to age matched controls (P<0.05).

**Table 1.** Physical and Metabolic Characteristics of Dunkin Hartley Guinea Pigs.

|  | **CONTROL** | **RAP** | **RAP+MET** |
| --- | --- | --- | --- |
| **Body Weight (g)** | 992 ± 90 | 846 ± 49* | 775 ± 100* |
| **Medial Tibia OARSI Score** | 4.6 ± 1.4 | 8.9 ± 3.0* | 8.7 ± 3.6* |
| **Fasted Blood Glucose (mg/dL)** | 242 ± 55 | 396 ± 61* | 335 ± 53* |
| **Blood Rapamycin (ng/mL)** | 0.4 ± 0 | 72 ± 8 | 78 ± 10 |
| **Blood Metformin (ng/mL)** | 2 ± 0 | N/A | 282 ± 54 |

**Note: Control values for Rap and Met in blood circulation are set to lowest readable value.**
**\*P<0.05 vs Con.**

### Mitochondrial Respiration

#### ADP bolus

When provided a saturating bolus of ADP there was no statistical differences in mitochondrial respiration between DH guinea pigs treated with Rap and Rap+Met compared to control (Figure 2A). There were also no differences between treatments compared to control in the P/E ratio (Figure 2B).

**Figure 1.**
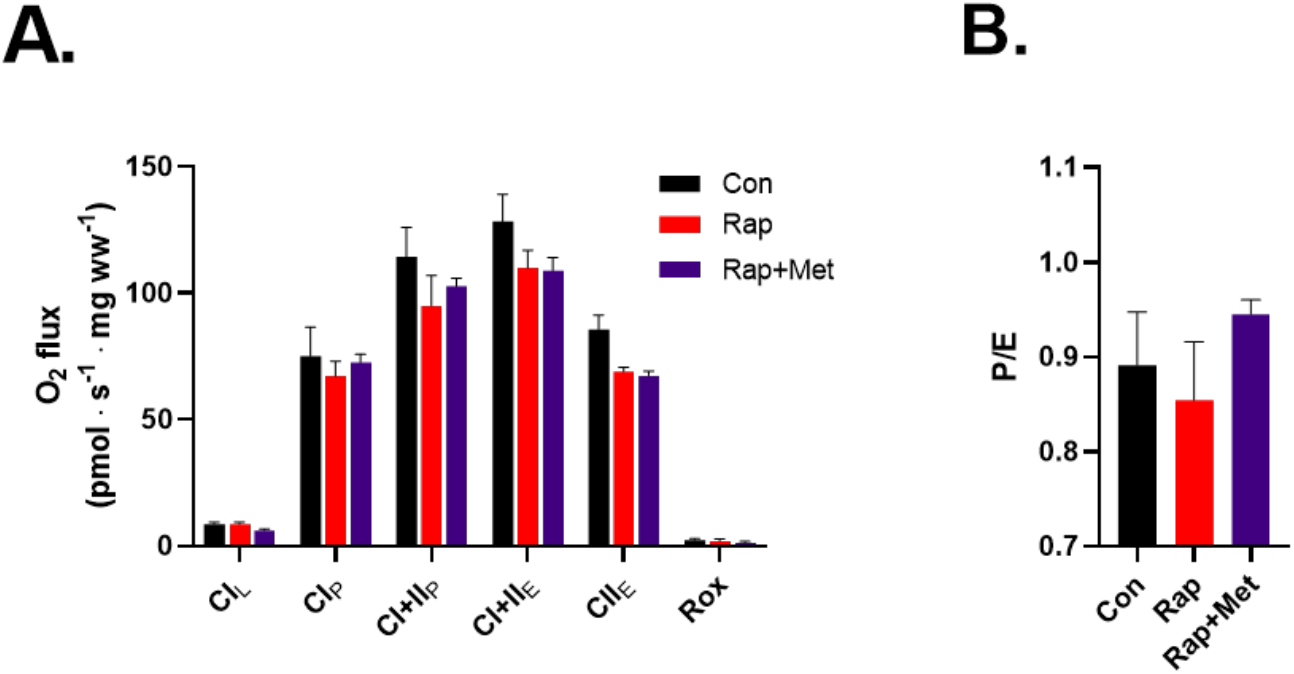
**A**) Skeletal muscle mitochondrial respiration during a SUIT protocol with an ADP bolus. **B**) The ratio of coupled OXPHOS to uncoupled ETS capacity (P/E). Con (n=4), Rap (n=4), and Rap+Met (n=3). Data are presented as mean ± SEM.

**Figure 2.**
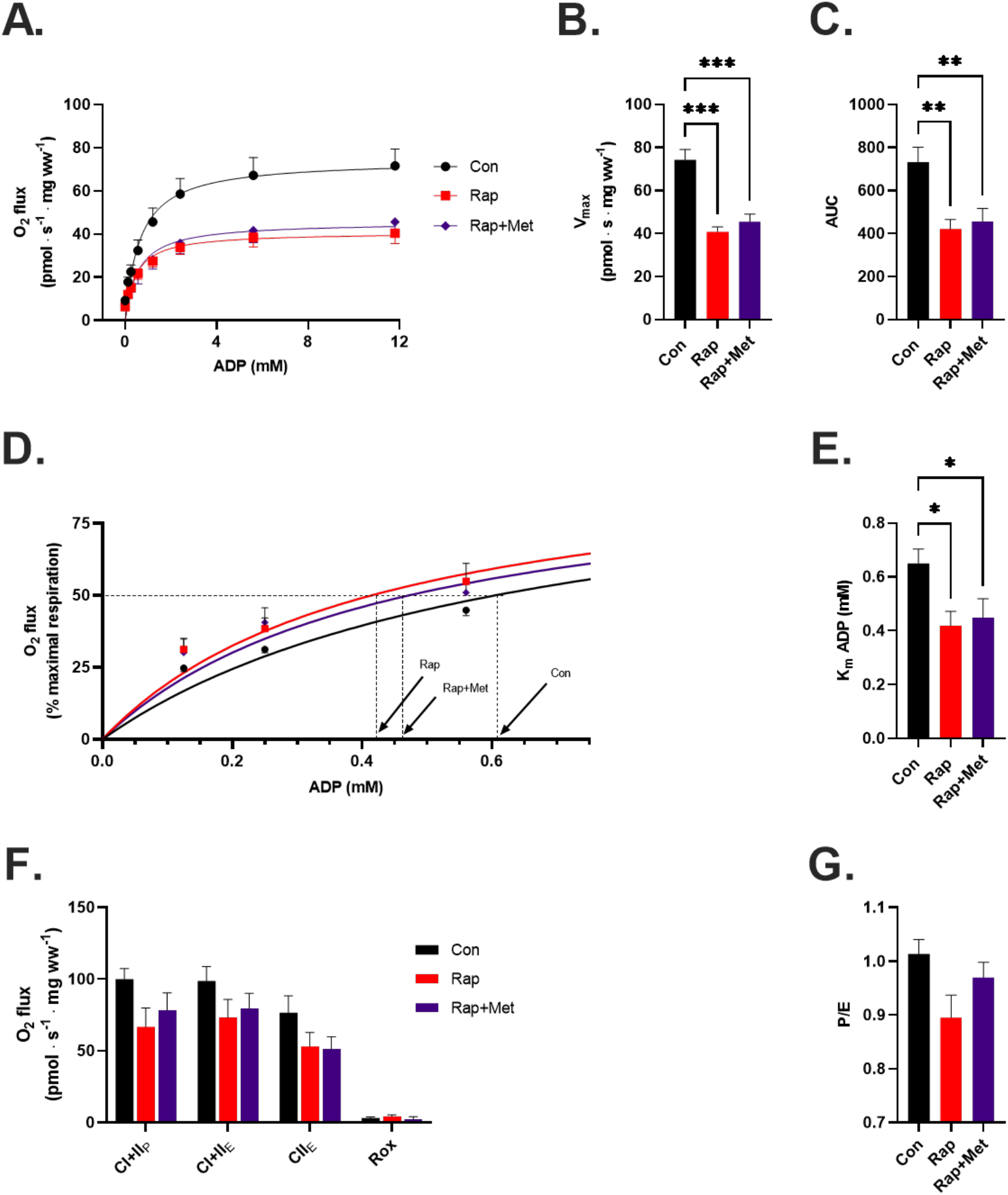
**A**) Complex I (CI)-linked respiration, **B**) Vmax and **C**) area under the curve (AUC) during an ADP titration protocol. **D**) CI-linked respiration expressed as a percentage of maximal respiration and used to determine **E**) the apparent Km of ADP. **F)** CI+II OXPHOS, ETS and ROX after the ADP titration protocol and **G)** the P/E ratio. Con (n=4), Rap (n=4), and Rap+Met (n=4). *P<0.05 vs. Con, **P<0.01 vs Con, ***P<0.001 vs Con. Data are presented as mean ± SEM

#### ADP Titration

Using the ADP titration protocol revealed statistically significant differences in DH guinea pigs treated with Rap and Rap+Met compared to control. Treatments did not influence CI_L_ respiration. Compared to control, Rap and Rap+Met treated DH guinea pigs had lower (P<0.05) submaximal and maximal CI-linked respiration starting at a sub-saturating dose of 1 mM ADP and continuing until a maximal, saturating dose of 12 mM ADP (Figure 3A). The inhibitory effects of Rap and Rap+Met were further supported by a lower V_max_ (P<0.001) and AUC (P<0.01) versus control (Figure 3B, C). Rapamycin also induced a leftward shift in the respiration curve (% of maximum) consistent with a lower apparent Km of ADP in the Rap and Rap+Met (P<0.05) treated DH guinea pigs compared to control (Figure 3D, E). A lower apparent Km of ADP represents a greater ADP sensitivity with Rap treatment. The differences in mitochondrial respiration between Rap and Rap+Met versus control were no longer apparent during CI+II_P_ (P=0.1), CI+II_E_, or CII_E_ (Figure 3F). Furthermore, the ratio of P/E was not significantly different after Rap or Rap+Met treatment (Figure 3G).

**Figure 3.**
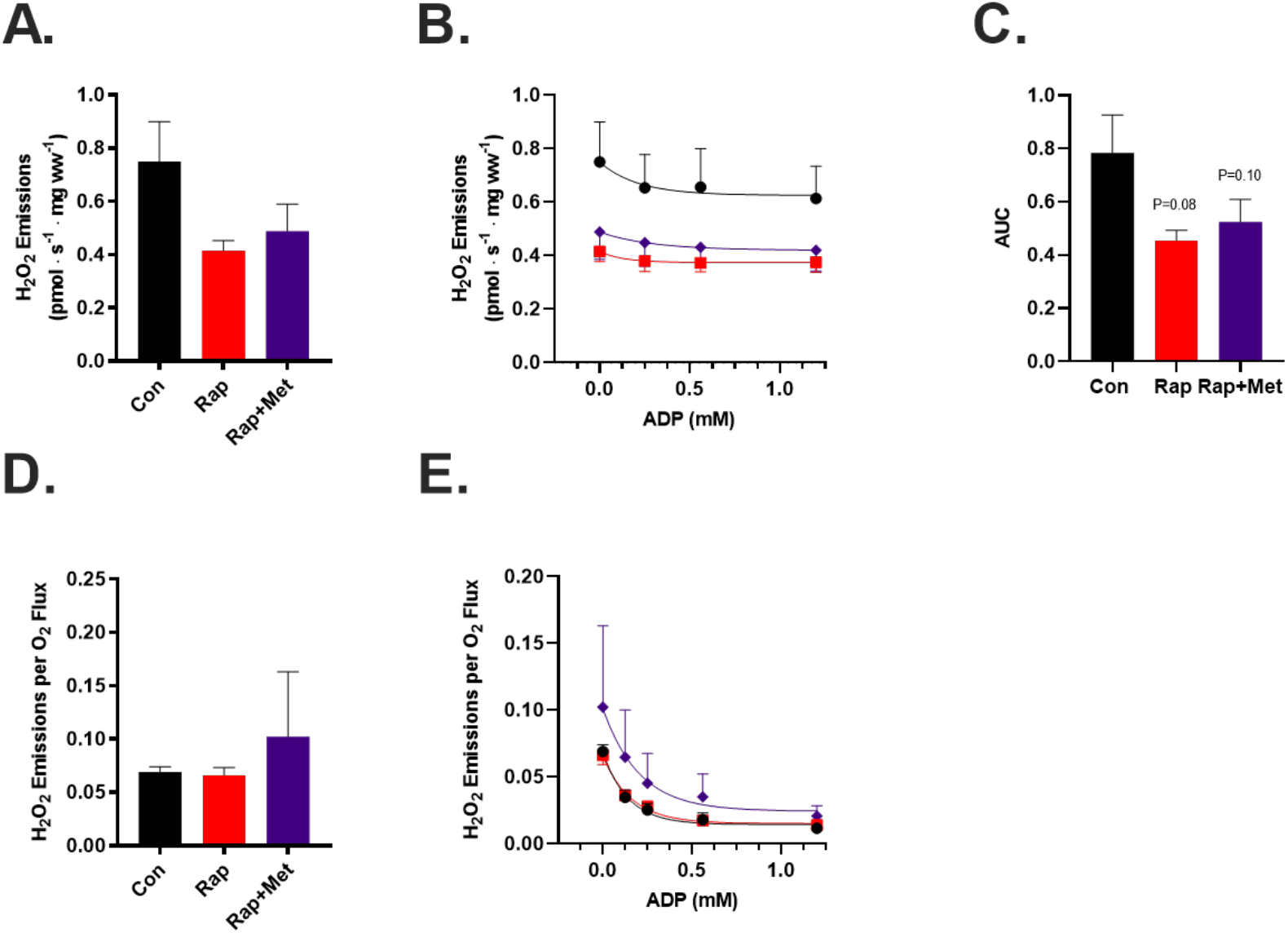
**A**) Skeletal muscle mitochondrial hydrogen peroxide (H_2_O_2_) emissions during CI-linked LEAK respiration and **B**) ADP titration. **C**) H_2_O_2_ emissions area under the ADP titration curve (AUC). **D**) Skeletal muscle H_2_O_2_ emissions expressed relative to respiration during CI-linked LEAK respiration and **E**) during titration of ADP. Con (n=4), Rap (n=4), and Rap+Met (n=4). Data are presented as mean ± SEM.

### Mitochondrial H_2_O_2_ emissions

Mitochondrial H_2_O_2_ emissions during CI-supported LEAK respiration tended to be lower in the Rap (P=0.11) and Rap+Met (P=0.12) treatments versus control (Figure 4A). This trend continued with titration of ADP as evident by a lower Mitochondrial H_2_O_2_ emissions AUC in guinea pigs treated with Rap (P=0.08) and Rap+Met (P=0.10) (Figure 4B, C). However, when mitochondrial H_2_O_2_ emissions were expressed relative to mitochondrial O_2_ flux, there were no longer trends for differences between groups (Figure 4D, E).

**Figure 4.**
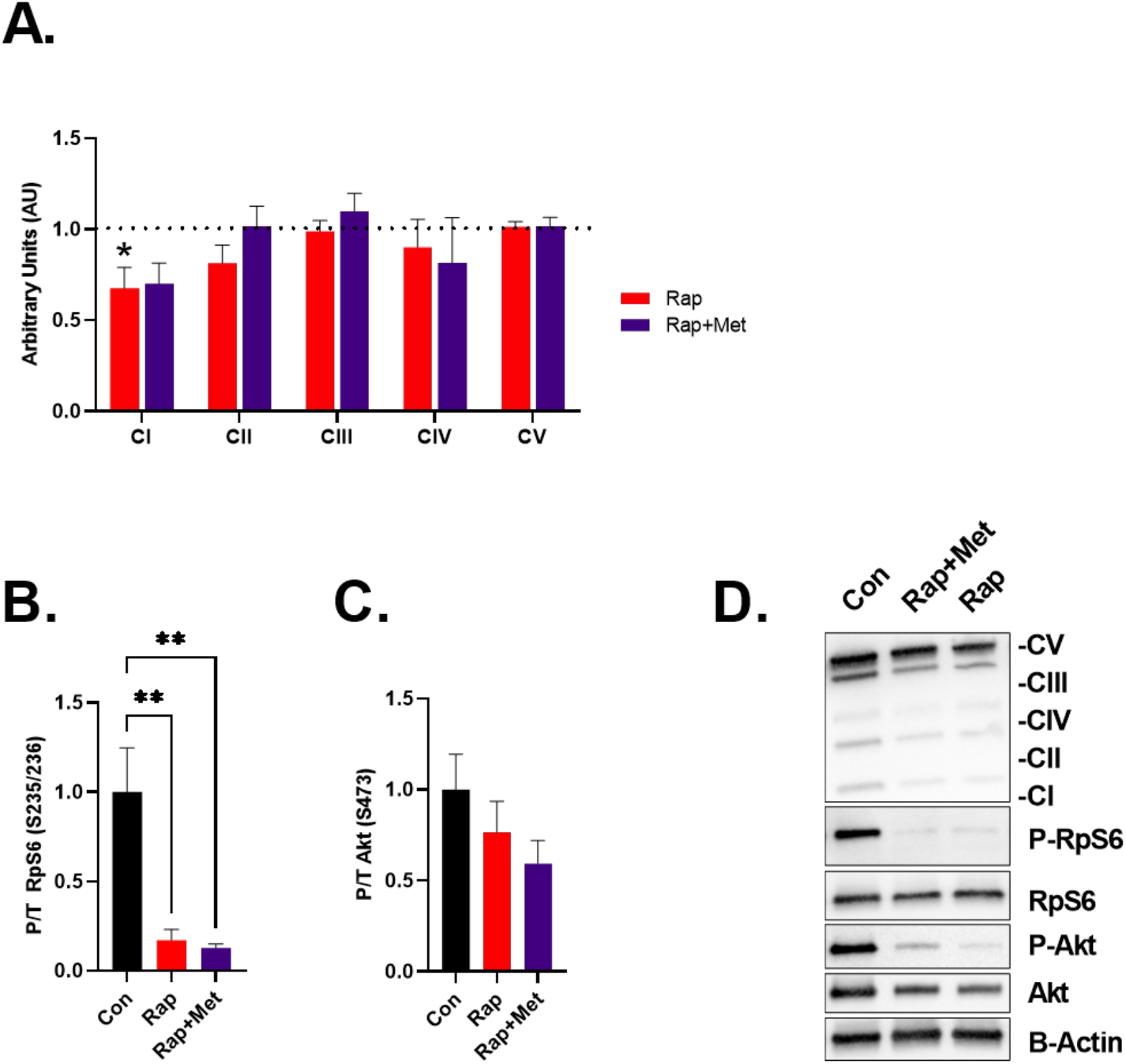
Skeletal muscle OXPHOS and nutrient-sensing signaling proteins. **A)** Protein content of the ETS complex I-V. Control is indicated by the dotted line and all data is expressed relative to the mean of the control. **B)** Protein content of P-RPS6/Total and **C)** P-AKT/Total. **D)** Protein content determined by western blot. Con (n=8), Rap (n=8), and Rap+Met (n=6).*P<0.05 vs. Con. **P<0.01 vs Con. Data are presented as mean ± SEM.

### Mitochondrial Content

To estimate mitochondrial protein abundance, we evaluated subunit content of complexes I through V of the ETS. CI protein content was lower (P<0.05) in the Rap and tended to be lower (P=0.09) in the Rap+Met treatment groups (Figure 4A). There were no other changes in protein abundance for any other complexes between DH guinea pigs receiving treatment compared to control.

### Nutrient Signaling

To evaluate whether dietary Rap and Rap+Met modified mTOR signaling pathways in skeletal muscle of the DH guinea pig, we investigated phosphorylation of RpS6 S235/236 and AKT S473 as downstream targets of mTOR complex 1 (mTORC1) and mTOR complex II (mTORC2). Rap and Rap+Met diminished mTORC1 signaling as evident by an 83 and 87% (P<0.01) decrease in phosphorylated RpS6 (Figure 4B) respectively. Rap and Rap+Met did not inhibit mTORC2 signaling as evident by a non-significant 20 and 40% decrease in AKT S473 phosphorylation (Figure 4C). For unknown reasons, we were unable to detect total or phosphorylated AMPK in skeletal muscle of DH guinea pigs. We therefore explored the impact of Rap and Rap+Met on AMPK signaling in the liver. While we were able to detect AMPK in the liver, no treatment altered AMPK phosphorylation (Data not shown).

## DISCUSSION

The main findings of this study are that at doses previously shown to slow aging in mice, dietary Rap and Rap+Met modified skeletal muscle mitochondrial bioenergetics in the DH guinea pig model of age-related primary OA. DH guinea pigs treated with Rap with or without Met had 1) lower submaximal and maximal CI-linked respiration, 2) greater ADP sensitivity, indicated by a lower apparent Km of ADP, and 3) less CI protein content. The rapamycin-induced changes in mitochondrial function recapitulate similar findings in human patients with severe, end-stage OA. Consistent with this notion, the impaired mitochondrial function in Rap treated DH guinea pigs were accompanied by elevated hyperglycemia and increased OA pathology. Collectively our data indicate that dietary Rap with or without Met appears to worsen skeletal muscle mitochondrial function, glycemic control, and OA pathology in DH guinea pigs.

### Rapamycin impairs mitochondrial function in a model of age-related OA

Interventions targeted to improve mitochondrial biogenesis and function have been proposed to increase longevity and delay musculoskeletal disease progression. Rap has demonstrated the ability to improve lifespan and extend healthspan in aging models (Harrison et al., 2009; Miller et al., 2011) but the effects on the mitochondria and in models of specific age-related disease are poorly understood. In the current study, we observed divergent findings depending on the protocol used to evaluate mitochondrial respiration in permeabilized muscle fibers. When using a bolus of ADP, there were no differences in maximal mitochondrial respiration between any treatment and control. These findings are not surprising and are in agreement with our previous study that showed by using a titration and not a bolus of ADP, we were able to detect subtle differences in skeletal muscle mitochondrial respiration after healthspan extending treatments in older adults (Konopka et al., 2019). Titrating ADP revealed that dietary Rap seemed to impair mitochondrial function in DH guinea pigs as evident by decreased submaximal and maximal CI-linked respiration and increased ADP sensitivity. These findings are in line with previous work to suggest that Rap decreased respiration in cultured muscle cells (Ye et al., 2012, 2013) and mice (Cunningham et al., 2007). In the context of aging, we initially viewed greater ADP sensitivity after Rap and Rap+Met treatment as beneficial since older adults have lower ADP sensitivity (Holloway et al., 2018). However, in patients with severe, end-stage OA about to undergo joint replacement, skeletal muscle mitochondrial ADP sensitivity was elevated and mitochondrial complex IV activity was lower compared to healthy, age-matched controls (Eimre et al., 2006; Safdar et al., 2010). We have previously found that the off-target side effects of dietary Rap and Rap+Met were associated with increased OA severity in DH guinea pigs. Therefore, our data in DH guinea pigs treated with Rap and Rap+Met are consistent with end-stage human OA and indicate a potential connection between greater OA severity and impaired skeletal muscle mitochondrial bioenergetics.

Alterations to adenylate transport proteins through either change in protein content or post-translational modifications contribute to the regulation of ADP sensitivity (Holloway et al., 2018) (Miotto et al., 2018). Transport of adenylates (ADP, ATP, etc.) into the mitochondria is mediated by voltage-dependent anion channels (VDAC), mitochondrial creatine kinase and adenine nucleotide transport proteins. Previous work indicates that mTORC1 phosphorylates Bcl-lx at serine 62 which complexes with VDAC1 to increase substrate and adenylate transport across the outer mitochondrial membrane (Ramanathan & Schreiber, 2009). Attenuation of mTORC1 signaling by Rap, as observed in the current study, can disassociate Bcl-lx from VDAC limiting ADP/ATP transport and could be one mechanism involved in lowering the apparent Km of ADP and limiting mitochondrial respiratory capacity.

### Rapamycin reduces CI abundance

The inhibition of CI-supported respiration after Rap treatment was accompanied by a decrease in CI protein abundance. The P/E ratio is considered a reflection of intrinsic mitochondrial function independent from changes in protein content. Therefore, no change in P/E ratio after Rap supports the notion that a decrease in CI content contributes to lower CI-linked respiration. Previous work has demonstrated that rapamycin decreases transcription factors involved in mitochondrial biogenesis (Cunningham et al., 2007; Ye et al., 2012, 2013) with no change in OXPHOS proteins. Rap preserves bulk mitochondrial protein synthesis rates (Drake et al., 2013) while proteomic approaches reveal that turnover of proteins within the ETS are subunit specific (Karunadharma et al., 2015; Wolff et al., 2020). Rap increased the half-life of the CI subunit NDFUB8 in skeletal and cardiac muscle (Dai et al., 2014; Karunadharma et al., 2015) which is the specific subunit probed in our western blot analysis for CI. Therefore, changes in NDUFB8 protein turnover could be one factor contributing to lower abundance and changes to mitochondrial respiration. Although it remains unclear how Rap alters mitochondrial content and function, it may be linked to changes in protein turnover and assembly of ETS subunits. Future work is needed to connect ETS proteostasis to functional outcomes to further understand the mechanisms promoting healthspan extension.

### Rapamycin decreases mitochondrial H_2_O_2_ emissions in a model of age-related OA

Mitochondrial H_2_O_2_ emissions in permeabilized muscle fibers are the result of H_2_O_2_ production and antioxidant scavenging capacity. Rap treatment tended to lower mitochondrial H_2_O_2_ emissions in the DH guinea pig. However, it should be noted that decreased mitochondrial H_2_O_2_ emissions with Rap treatment were driven primarily in the absence of ADP during CI-linked LEAK respiration. Future studies should consider also using substrates with convergent electron supply to the Q-junction through CII to induce maximal mitochondrial H_2_O_2_ emissions to clarify if Rap also improves the sensitivity of ADP to suppress H_2_O_2_ emissions. When mitochondrial H_2_O_2_ emissions were expressed relative to respiration, there were no longer differences between Rap and control. The ratio of mitochondrial H_2_O_2_ emissions to respiration is often an index to reflect electron leak. Since Rap similarly decreases both respiration and H_2_O_2_ emissions, these data would suggest that Rap may influence flux through OXPHOS.

### Elevated plasma glucose after 12 weeks of rapamycin treatment

A common side effect of chronic rapamycin treatment is impaired glucose metabolism and insulin resistance driven by off target inhibition of mTORC2 (Arriola Apelo, Pumper, et al., 2016; Lamming et al., 2012; Ye et al., 2012). Consistent with these findings, we show that 12-weeks of dietary Rap and Rap+Met increased plasma glucose and non-significantly attenuated skeletal muscle mTORC2 signaling as evident by a 20% and 40% decrease in AKT S473 phosphorylation. We have also found that increased plasma glucose was correlated to increased OA severity in DH guinea pigs. Intermittent rapamycin dosing schedules or alternative rapamycin analogs that selectively inhibit mTORC1 minimize off-target side metabolic effects mediated by inhibition of mTORC2 (Arriola Apelo, Neuman, et al., 2016). Therefore, future work is needed to understand if different rapamycin or rapalog dosing regimens may prevent detrimental side effects such as impaired mitochondrial function and hyperglycemia and be able to delay or prevent age-related OA pathology.

## Conclusion

Within the current study we observed that DH guinea pigs treated with Rap either alone or in combination with Met, had lower mitochondrial CI-linked respiration and greater ADP sensitivity. These findings are consistent with previous work in patients with end-stage OA that have lower mitochondrial respiration and elevated ADP sensitivity compared to age-matched non-OA controls. Taken together with our previous findings that Rap-induced hyperglycemia was associated with greater OA severity, our data suggests long-term treatment with dietary rapamycin (14ppm) may not be suitable for the prevention or treatment of age-related OA in DH guinea pigs. Additional work is needed to determine if alternative rapamycin dosing regimens such as lower concentrations, intermittent administration, or rapamycin analogs may be more effective to safely maximize healthspan extending effects on OA pathology by minimizing off-target side effects associated with chronic use of dietary rapamycin.

